# An extensive enhancer-promoter map generated by genome-scale analysis of enhancer and gene activity patterns

**DOI:** 10.1101/190231

**Authors:** Tom Aharon Hait, David Amar, Ron Shamir, Ran Elkon

## Abstract

Massive efforts have documented hundreds of thousands of putative enhancers in the human genome. A pressing genomic challenge is to identify which of these enhancers are functional and map them to the genes they regulate. We developed a novel method for inferring enhancer-promoter (E-P) links based on correlated activity patterns across many samples. Our method, called *FOCS*, uses rigorous statistical validation tailored for zero-inflated data, identifying the most important E-P links in each gene model. We applied FOCS to the wide epigenomic and transcriptomic datasets recorded by the *ENCODE*, *Roadmap Epigenomics* and *FANTOM5* projects, together covering 2,630 samples of human primary cells, tissues and cell lines. In addition, building on expression of enhancer RNAs (eRNAs) as an exquisite mark of enhancer activity and on the robust detection of eRNAs by the GRO-seq technique, we compiled a compendium of eRNA and gene expression profiles based on public GRO-seq data from 245 samples and 23 human cell types. Applying FOCS to this compendium further expanded the coverage of our inferred E-P map. Benchmarking against gold standard E-P links from ChIA-PET and eQTL data, we demonstrate that FOCS prediction of E-P links outperforms extant methods. Collectively, we inferred >300,000 cross-validated E-P links spanning ^~^16K known genes. Our study presents an improved method for inferring regulatory links between enhancers and promoters, and provides an extensive resource of E-P maps that could greatly assist the functional interpretation of the noncoding regulatory genome. FOCS and our predicted E-P map are publicly available at http://acgt.cs.tau.ac.il/focs.

## Introduction

Deciphering the regulatory role of the noncoding part of the human genome is a major challenge. With the completion of the sequencing of the genome, efforts have shifted over the last decade towards understating the epigenome. These efforts aim at understanding regulatory mechanisms outside the protein-coding sequences that allow the production of thousands of different cell types from the same DNA blueprint. Enhancer elements that distally control the activity of target promoters play critical roles in this process. Consequently, large-scale epigenomic projects set out to identify all the cis-regulatory elements that are encoded in the genome. Prominent among them is the ENCODE consortium [1,2], which applied a variety of epigenomics techniques to a large panel of human cell lines. Profiling epigenetic marks of regulatory activity (including DHS-seq profiling of DNase I hypersensitive sites, which is accepted as a common feature of all active elements), ENCODE collectively identified hundreds of thousands of putative regulatory elements in the genome [2]. As ENCODE analyses were mainly applied to cancer cell lines, a follow-up project, the Roadmap Epigenomics, applied similar analyses to a large collection of human primary cells and tissues, in order to establish more physiological maps of common and cell-type specific putative regulatory elements [3]. Given the plethora of candidate enhancer regions called by these projects, the next pressing challenge is to identify which of them is actually functional and map them to the genes they regulate. A naïve approach that is still widely used in genomic studies links enhancers to their nearest genes. Yet, emerging indications suggest that up to 50% of enhancers cross over their most proximal gene and control a more distal one [4]. A common approach that improves this naïve E-P mapping is by using pairwise correlation, which calculates activity patterns for each promoter (P) and putative enhancer (E) over the probed cell panel, and identifies E-P pairs, located within a distance limit, that show highly correlated patterns across many samples [2,3]. However, this approach does not take into account interactions among multiple enhancers that control the same target promoter. Furthermore, Pearson correlation, which is typically applied for this task, is highly sensitive to outliers and thus prone to false positives.

Improved detection of functional enhancers is offered by a recently discovered class of non-coding transcripts, named enhancer RNAs (eRNAs) [5]. eRNAs are mostly transcribed bi-directionally from regions of enhancers that are actively engaged in transcriptional regulation [5] (reviewed in [6,7]), and, importantly, changes in eRNA expression at specific enhancer regions in response to different stimuli correlate both with changes in the amount of epigenetic marks at these enhancers and with the expression of their target genes [8–11]. Most eRNAs are not polyadenylated and are typically expressed at low levels due to their instability (reviewed in [12]). Therefore, eRNAs are not readily detected by standard RNA-seq protocols, but can be effectively measured by global run-on sequencing (GRO-seq), a technique that measures production rates of all nascent RNAs in a cell [8–10,13,14] or by cap-analysis of gene expression (CAGE) followed by sequencing [4,15,16]. Utilizing eRNA expression as a mark of enhancer activity, the FANTOM5 consortium recently generated an atlas of predicted enhancers in a large panel of human cancer and primary cell lines and tissues [4]. This study too used pairwise correlation (in this case, calculated between expression levels of an eRNA and a gene whose TSS is within a distance limit from it) to infer E-P links.Regression analysis was applied to characterize the architecture of promoter regulation by enhancers [4]. However, since all samples were used for training the regression models, this analysis is prone to over-fitting and thus their predictive power on new samples is unclear.

Here, we present *FOCS* (**F**DR-corrected **O**LS with **C**ross-validation and Shrinkage) - a novel procedure for inference of E-P links based on correlated activity patterns across many samples from heterogeneous sources. FOCS uses a cross-validation scheme in which regression models are learnt on a training set of samples and then evaluated on left-out samples from other cell types. The models are subjected to a new statistical validation scheme that is tailored for zero-inflated data. Finally, validated models are optimally reduced to derive the most important E-P links. We applied FOCS on massive genomic datasets recorded by ENCODE, Roadmap Epigenomics and FANTOM5, and on a large compendium of eRNA and gene expression profiles that we compiled from publicly available GRO-seq datasets. We demonstrate that FOCS outperforms extant methods in terms of concordance with E-P interactions identified by ChIA-PET and eQTL data. Collectively, applying FOCS to these four data resources, we inferred ^~^300,000 cross-validated E-P interactions spanning ^~^16K known genes. FOCS and our predicted E-P maps are publicly available at http://acgt.cs.tau.ac.il/focs.

## Results

### The FOCS procedure for predicting enhancer-promoter links

We set out to develop an improved statistical framework for prediction of E-P links based on their correlated activity patterns measured over many cell types. As a test case, we first focused on ENCODE's DHS profiles [2], which constitute 208 samples measured in 106 different cell lines (Online Methods) [2]. This rich resource was previously used to infer E-P links based on pairwise correlation between DHS patterns of promoters and enhancers located within a distance of ±500 kbp. Out of ^~^42M pairwise comparisons, ^~^1.6M pairs showed Pearson's correlation>0.7 and were regarded as putatively functional E-P links [2]. However, Pearson's correlation is sensitive to outliers and thus may be prone to high rate of false positive predictions. This is especially exacerbated in cases of sparse data (zero inflation), which are prevalent in enhancer activity patterns, as many of the enhancers will be active only in a limited set of conditions. In addition, the combinatorial nature of transcriptional regulation in which a promoter is regulated by multiple enhancers is not considered by such pairwise approach.

To address these points we developed a novel statistically-controlled regression analysis scheme for E-P mapping, that we dubbed FOCS. Specifically, FOCS uses regression analysis to learn predictive models for promoter's activity from the activity levels of its *k* closest enhancers, located within a window of ±500 kb around the gene's TSS. (Throughout our analyses we used *k =* 10). Importantly, to avoid overfitting of the regression models to the training samples, FOCS implements a leave-cell-type-out cross validation (LCTO CV) procedure, as follows. In a dataset that contains samples from *C* different cell-types, for each promoter, FOCS performs *C* iterations of model learning. In each iteration, all samples belonging to one cell-type are left out and the model is trained on the remaining samples. The trained model is then used to predict promoter activity in the left-out samples (**Fig. 1**).

**Figure 1.**
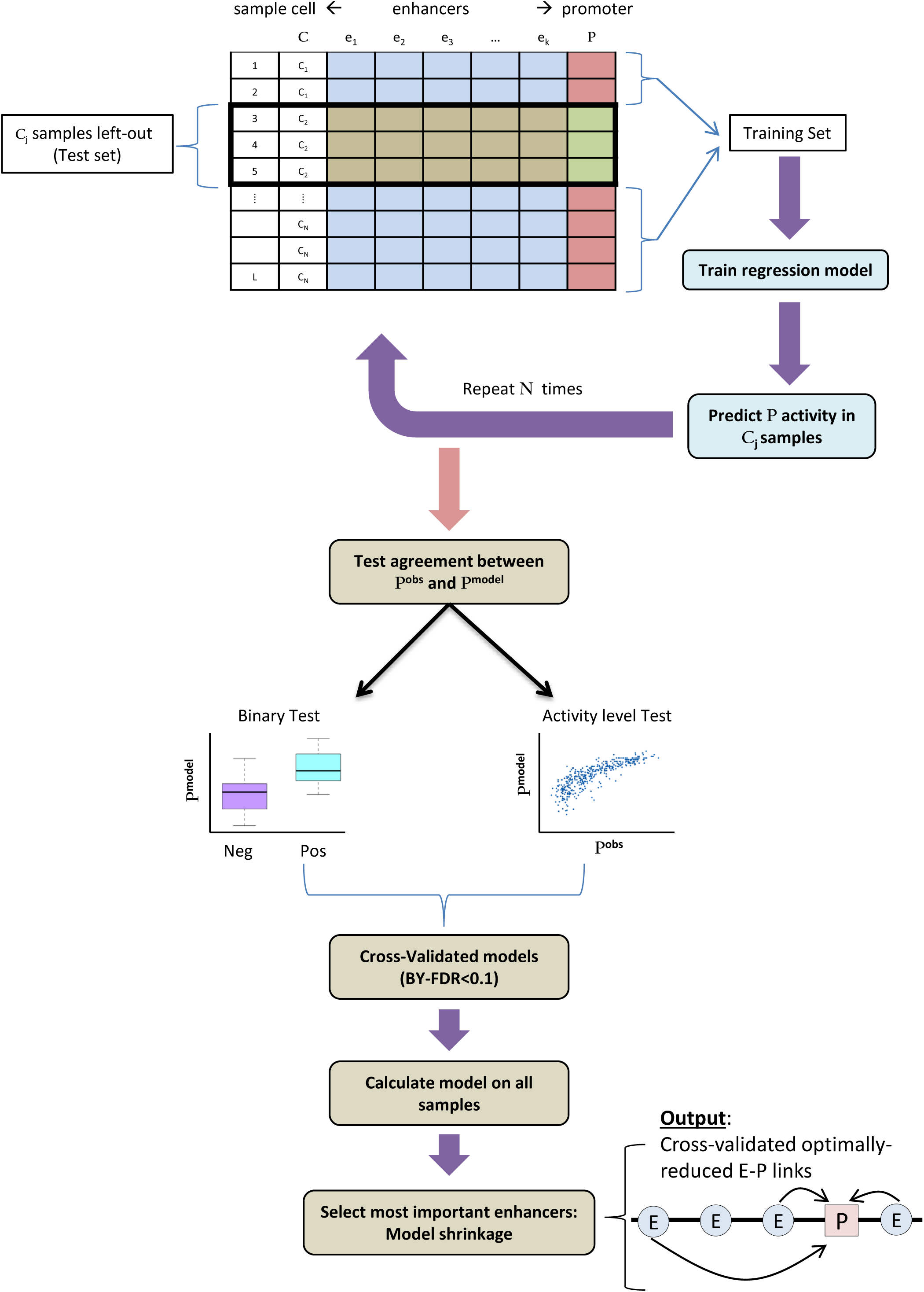
FOCS statistical procedure for inference of E-P links. In a dataset with samples from N different cell types, FOCS starts by performing N cycles of leave-cell-type out cross-validation (LCTO CV). In cycle *j*, the set of samples from cell-type *C*_*j*_ is left out as a test set, and a regression model is trained, based on the remaining samples, to estimate the level of the promoter P (the independent variable) from the levels of its k closest enhancers (the dependent variables). The model is then used to predict promoter activity in the test set samples. After the N cycles, FOCS tests the agreement between the predicted (P^model^) and observed (P^obs^) promoter activities using two non-parametric tests. In the *binary test*, samples are divided into positive (P^obs^ ≥1RPKM) and negative (P^obs^ <1RPKM) sets, and the ability of the inferred models to separate between the sets is examined using Wilcoxon rank-sum test. In the *activity level test*, the consistency between predicted and observed activities in the positive set of samples is tested using Spearman correlation. P-values are corrected using the BY-FDR procedure, and promoters that passed the validation tests (FDR≤0.1) are considered validated, and full regression models, this time based on all samples, are calculated for them. In the last step, FOCS shrinks each promoter model using elastic net to select its most important enhancers.

We implemented and evaluated three alternative regression methods: ordinary least squares (*OLS*), generalized linear model with the negative binomial distribution (*GLM.NB*) [17] and zero-inflated negative binomial (*ZINB*) [18]. GLM.NB accounts for unequal mean-variance relationship within subpopulations of replicates. ZINB is similar to GLM-NB but also accounts for excess of samples with zero entries (**Online Methods**). For each promoter and regression method, the learning phase yields an activity vector, containing the promoter's activity in each sample as predicted when it was left out. FOCS applies two non-parametric tests, tailored for zero-inflated data, to evaluate the ability of the inferred models (consisting of the k nearest enhancers) to predict the activity of the target promoter in the left-out samples. The first test is a "*binary test*" in which samples are divided into two sets, positive and negative, containing the samples in which the promoter was active or not, respectively, based on their measured signal. (We used a signal threshold of 1.0 RPKM for this classification.) Then, Wilcoxon signed-rank test is used to compare the predicted promoter activities between these two sets (**Fig. 1**). The second test is an "*activity level test*", which examines the agreement between the predicted and observed promoter's activities using Spearman's correlation. In this test, only the positive samples (that is, samples in which the measured promoter signal is ≥1.0 RPKM) are considered. Gene models with good predictive power should discriminate well between positive and negative samples (the binary test) and preserve the original activity ranks of the positive samples (the activity level test), and models that pass these tests are regarded as statistically cross-validated. Of note, these two validation tests evaluate each promoter model non-parametrically without assuming any underlying distribution on the data when inferring significance. Next, FOCS corrects the p-values obtained by these tests for multiple testing using the Benjamini and Yekutieli (BY) FDR procedure [19] with q-value<0.1. The BY FDR procedure takes into account possible positive dependencies between tests while the more frequently used Benjamini and Hochberg (BH) FDR procedure [20] assumes the tests are independent.

### FOCS results for ENCODE DHS epigenomic data

Applying FOCS to the ENCODE DHS dataset, we only considered promoters and enhancers that were active (that is, with signal > 1.0 RPKM) in at least 30 out of the 208 samples. Overall, this dataset contained 92,909 and 408,802 active promoters and enhancers, respectively (**Online Methods**). We first evaluated the performance of the three alternative regression methods in terms of the number of validated models each of them yielded. We found that the OLS method consistently produced more validated models that passed both the binary and activity level tests (**Fig. 2A-B; Supplementary Table 1**). Out of the 92,909 analyzed promoters, 52,658 had models that passed both tests (q-value≤0.1), while for 7,007 promoters, models passed none of these two tests (**Fig. 2C**). As expected, promoters with models that passed only the activity level test were active in a very high number of samples while those with models that passed only the binary test were active in much lower number of samples (**Fig. 2D**) (see **Supplementary Fig. 1** for examples of promoters in different validation categories). To examine the effect of the leave-cell-type-out cross validation (CV) procedure we compared *R*^2^ values obtained by OLS models generated without CV to the values obtained when CV was applied (**Fig. 2E**). The results indicate that without CV, many models are over-fitted to the training samples and have low predictive power on new ones. This problem is more severe in other datasets that we analyzed, as shown in subsequent section. **Fig. 2F** shows an example of promoter model with low predictive power on new samples, and demonstrates the high sensitivity of Pearson's correlation (or equivalently, of R^2^) to outliers. Such promoter models do not pass our CV tests and are considered to have low confidence.

**Figure 2.**
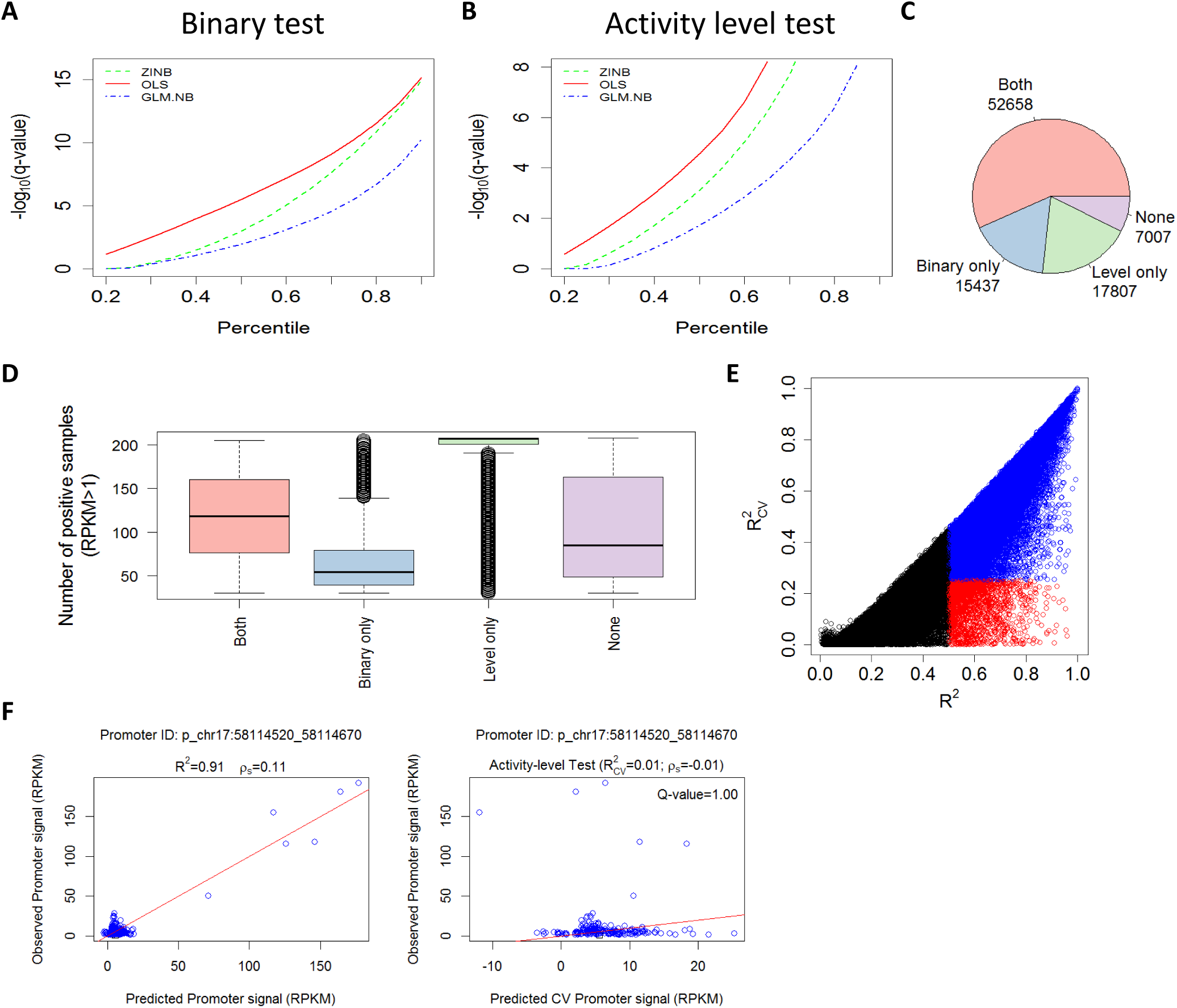
Performance of three alternative regression methods for inferring E-P models. (a) Performance of optimal least squares (OLS), generalized linear model with negative binomial distribution (*GLM.NB*) and zero-inflated negative binomial (*ZINB*) regression using the binary test. Point (x,y) on a plot indicates that a fraction x of the models had ‐log_10_[q-values] < y computed by Wilcoxon rank sum test. OLS yields a higher fraction of validated models at any q-value cutoff. (b) Same as (a) but using the activity level validation test, with p-values computed by the Spearman correlation test. Here too, OLS yields a higher fraction of validated models than the other methods. (c) Number of promoters whose OLS models passed (at q<0.1) each of the tests (or none). (d) The distribution of the number of positive samples (samples in which the promoter is active, i.e., has RPKM ≥ 1) for promoters in each category. (e) Comparison between the *R*^2^ values with/without cross-validation (CV). Each dot is a promoter model. Blue dots denote models with *R*^2^ ≥ 0.5 and 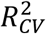 ≥ 0.25. Red dots denote models with and *R*^2^ ≥ 0.5 and 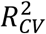 < 0.25 corresponding to over-fitted models with low predictive power on novel samples. (f) A promoter whose model as computed without CV gets very high *R*^2^ (left plot) but when CV is applied a low 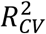 is obtained (right plot). This example demonstrates the sensitivity of *R*^2^ (and Pearson correlation) to outliers. *ρ*_*s*_: Spearman correlation, Q-value: FDR corrected P-value.

### The architecture of promoter regulation by enhancers

Next, we sought to characterize the architecture of promoter regulation by its enhancers, in terms of the number of regulating enhancers and their relative contribution. For each promoter that passed the validation tests, we now calculated a final model, this time considering all samples (**Fig. 1**), and estimated the relative contribution of each of its k enhancers to this full model. As in [4], per model, we measured the proportional contribution of each enhancer by calculating the ratio *r*^2^/*R*^2^ where *r* is the pairwise Pearson correlation between the enhancer and promoter activity patterns and *R*^2^ is the coefficient of determination of the entire promoter's model. In the analysis of the ENCODE DHS data, we included in this step the 70,465 promoters that passed the activity level test (or both tests). In agreement with previous observation [4], the closest enhancers have significantly higher contribution than the distal ones (**Fig. 3A**). However, the proportional contribution quickly reaches a plateau, indicating that above a certain threshold, distance to promoter is no longer an important factor, and enhancers #6-#10 (ordered according to their distance from the promoter) contribute similarly to promoter activity (**Fig. 3A**). Second, we examined the distribution of *R*^2^ values of these statistically validated models. 54% of the models (37,716 out of 70,465) had *R*^2^ ≥ 0.5 (**Fig. 3B**). 61% of the 52,658 models that passed both tests had *R*^2^ ≥ 0.5, compared to 32% of the 17,807 models that passed only the activity level test. (In contrast, only 13% of 15,437 models that passed only the binary test had *R*^2^ ≥ 0.5). We note that models that passed the CV tests but have low *R*^2^ do contain confident and predictive information on E-P links, though the low *R*^2^ suggests that there are additional missing regulatory elements that play important roles in the regulation of the target promoter.

**Figure 3.**
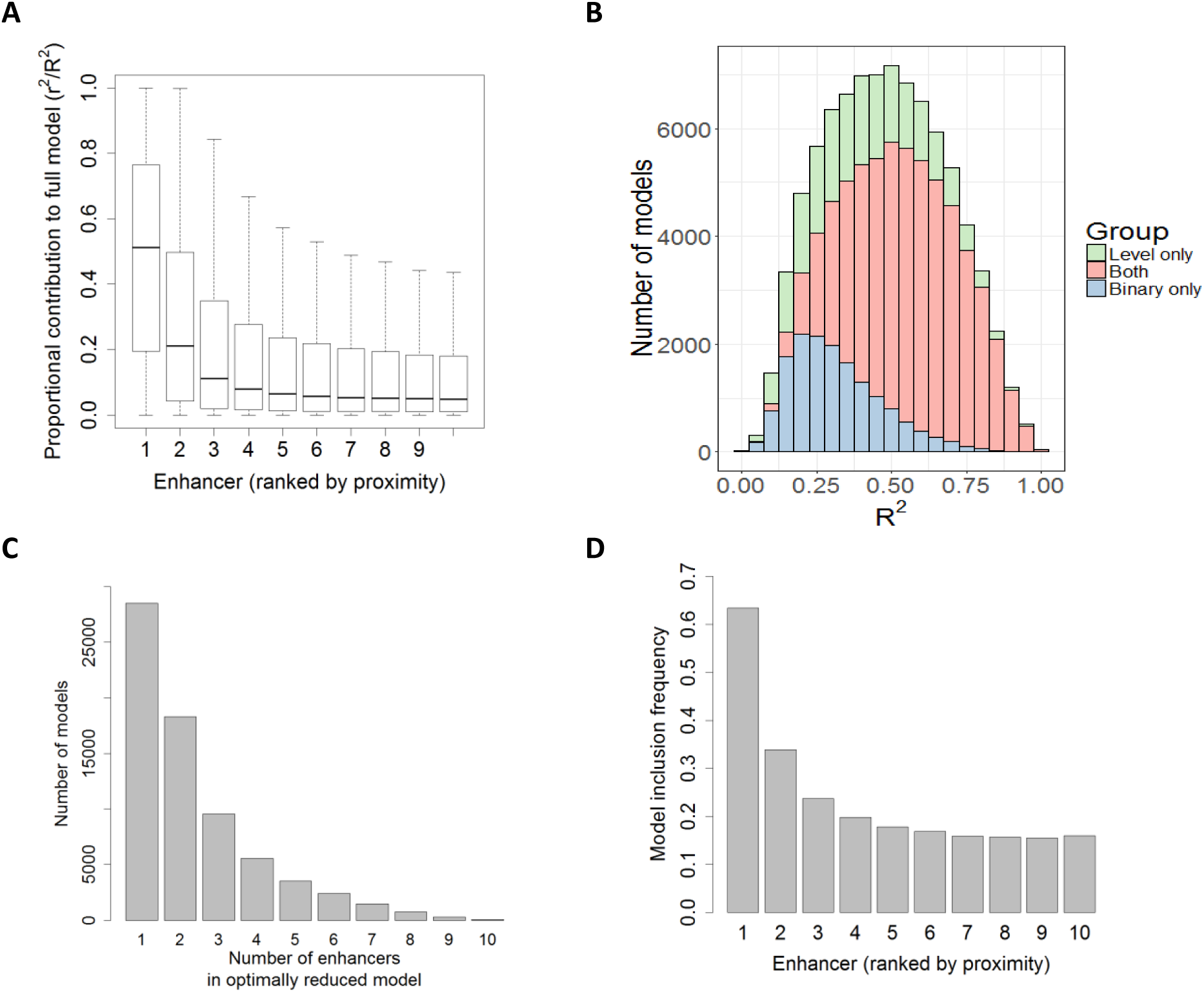
Architecture of promoter regulation by enhancers. (a) The proportional contribution of the 10 most proximal enhancers (within ±500kb of the target promoter) to models predicting promoter activity. The X axis indicates the order of the enhancers by their relative distance from the promoter, with 1 being the closest. (b) *R*^2^ values of the models that passed one or both CV tests. (c) Distribution of the number of enhancers included in the validated, optimally reduced models (i.e. after elastic net shrinkage). Most shrunken models contain 1-3 enhancers. (d) Inclusion rate of enhancers in the shrunken models as a function of their relative proximity to the target promoter.

A promoter's model produced by OLS regression contains all k variables (i.e., enhancers), where each variable is assigned a significance level (p-value) reflecting its statistical strength. Next, to focus on the most informative E-P interactions, FOCS seeks the strongest enhancers in each model. To this end, FOCS derives, per promoter, an optimally reduced model by applying model shrinkage (**Online Methods**). Lasso-based shrinkage was previously used for such task [4]. Here, we chose elastic-net *(enet)* approach, which combines Lasso and Ridge regularizations, since in cases of highly correlated variables (i.e., the enhancers), Lasso tends to select a single variable while Ridge gives them more equal coefficients (**Online Methods**). In this analysis too, we included the 70,465 models that passed the activity level test. **Fig. 3C** shows the distribution of the number of enhancers that were included in the enet-reduced models. On average, each promoter was linked to 2.4 enhancers. Inclusion rate decreased with E-P distance: the most proximal enhancer was included in 63% of the models while the 10^th^ enhancer was included in only 16% of them (**Fig. 3D**). Here too, the graph reaches a plateau and enhancers #6-#10 show very similar inclusion rates.

### Comparison of FOCS and extant methods performance using external validation resources

After optimally reducing the promoter models FOCS predicted in the ENCODE DHS dataset a total of 167,988 E-P links covering 70,465 promoters and 92,603 distinct enhancers (http://acgt.cs.tau.ac.il/focs/data/encode_interactions.txt). Next, we compared the performance of FOCS and three alternative methods for E-P mapping: (1) *Pairwise:* pairwise Pearson correlation > 0.7 between E-P pairs located within ±500 kbp, and accounting for multiple testing using BH (FDR <10^‒5^) (this was the main method used in [4], and also in [2] without multiple testing correction) (2) *OLS+LASSO:* Models are derived by OLS analysis using *all* samples without CV, selected based on *R*^2^ ≥ 0.5 and reduced using LASSO shrinkage (**Online Methods**) (this method was also applied in [4]). (3) *OLS+enet:* Same as (2) but with enet shrinkage in place of LASSO. **Table 1** summarizes the number of E-P links obtained by each method. FOCS yielded ^~^75% more models than the other methods.

**Table 1.** Number of inferred promoter models obtained by four alternative methods on the ENCODE DHS dataset

| Method type | #promoter models | #E-P links | #Unique enhancers |
| --- | --- | --- | --- |
| **Pairwise ($r \geq 0.7$)+ *FDR*** | 39,372 | 139,170 | 53,950 |
| **OLS-LASSO ($R^2 \geq 0.5$ )\*** | 39,368 | 122,064 | 74,104 |
| **OLS-enet ($R^2 \geq 0.5$ )\*** | 39,407 | 150,158 | 85,926 |
| **FOCS** | 70,465 | 167,988 | 92,603 |

(\*) The number of OLS models ( $R^2 \geq 0.5$ ) was 39,892 before LASSO / enet shrinkage. These methods eliminate models in which no enhancer passed the shrinkage.

To evaluate the validity of E-P mappings predicted by each method, we used two external omics resources: physical E-P interactions derived from ChIA-PET data and functional E-P links indicated by eQTL analysis. For E-P physical interactions, we used public ChIA-PET data that used RNAPII as the immunoprecipitated factor in MCF7, HCT-116, K562 and HelaS3 cell lines (a total of 922,997 interactions downloaded from the CCSI DB [21]). eQTL data was downloaded from the GTEx project (2,283,827 unique significant eQTL-gene pairs) [22]. We defined a 1 kbp interval for each promoter and enhancer and calculated the fraction of E-P links that were supported by either ChIA-PET or eQTL data (**Online Methods**). Remarkably, FOCS not only yielded many more E-P links (15,000-40,000 more), but also outperformed the alternative methods in terms of the fraction of predictions supported by either ChIA-PET (**Fig. 4A**) or eQTL data (**Fig. 4B**). **Figure 5** shows two FOCS-derived promoter models that are supported by ChIA-PET and eQTLs. Note that in CD4 promoter model (**Fig. 5B**) the 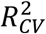 value was low (^~^0.1) while the Spearman correlation (*ρ*_*s*_) was 0.53 after CV. This demonstrates that FOCS can capture promoter models that exhibit non-linear relationship between the promoter and enhancer activities.

**Figure 4.**
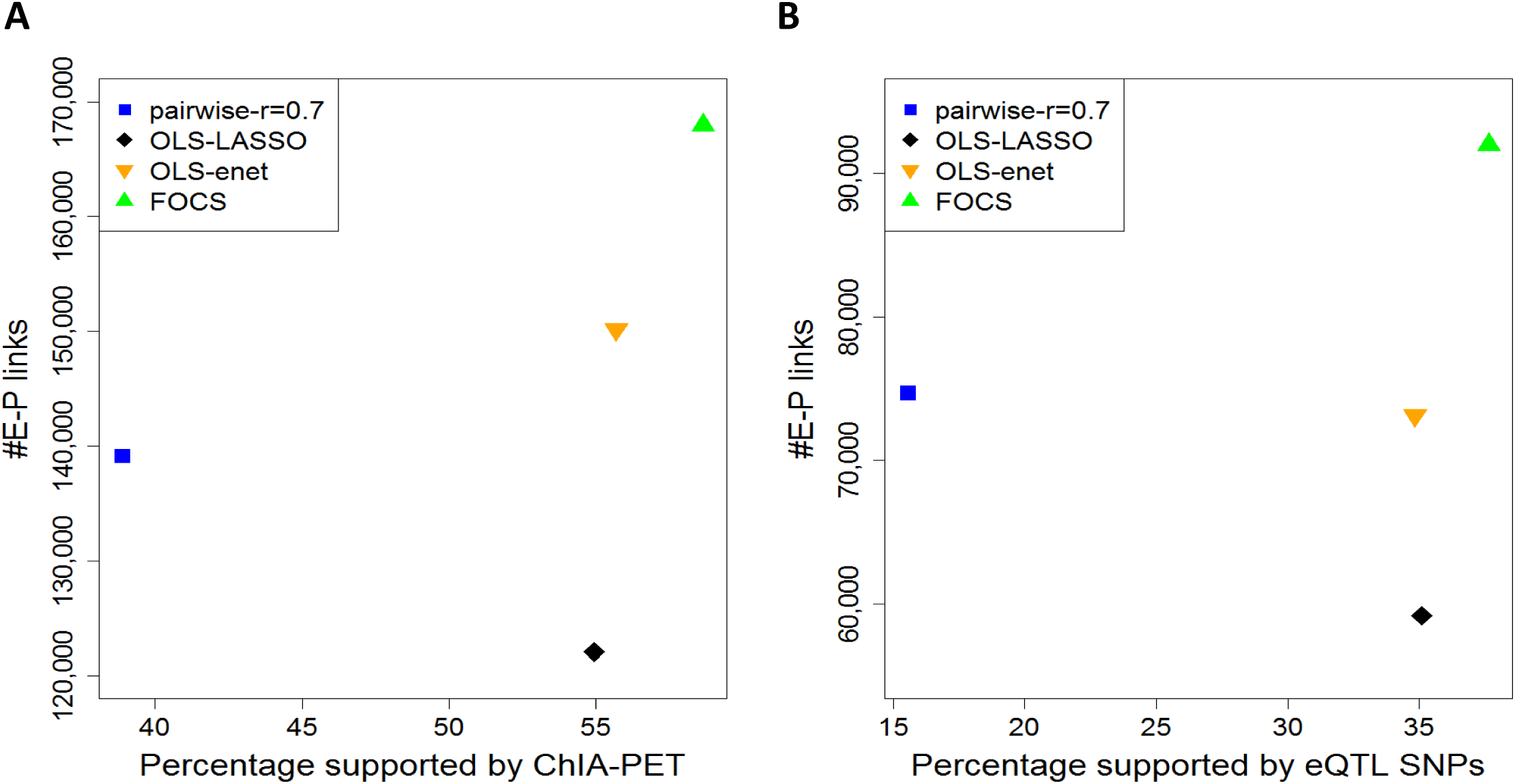
Comparison of the performance of different methods for predicting E-P links using ChIA-PET and eQTL data as external validation. Y-axis shows the total number of predicted E-P links. X-axis shows the percentage supported by the external source: (A) Pol-II ChIA-PET. (B) GTEX eQTLs. In (B) the y-axis shows the total number of predicted E-P links where the promoter is annotated with a known gene. FOCS (green triangle) makes more predictions and also manifests highest support rate by both ChIA-PET (59%) and eQTL (38%). In all methods, empirical p-value by random permutation test was < 0.01 (Online Methods).

**Figure 5.**
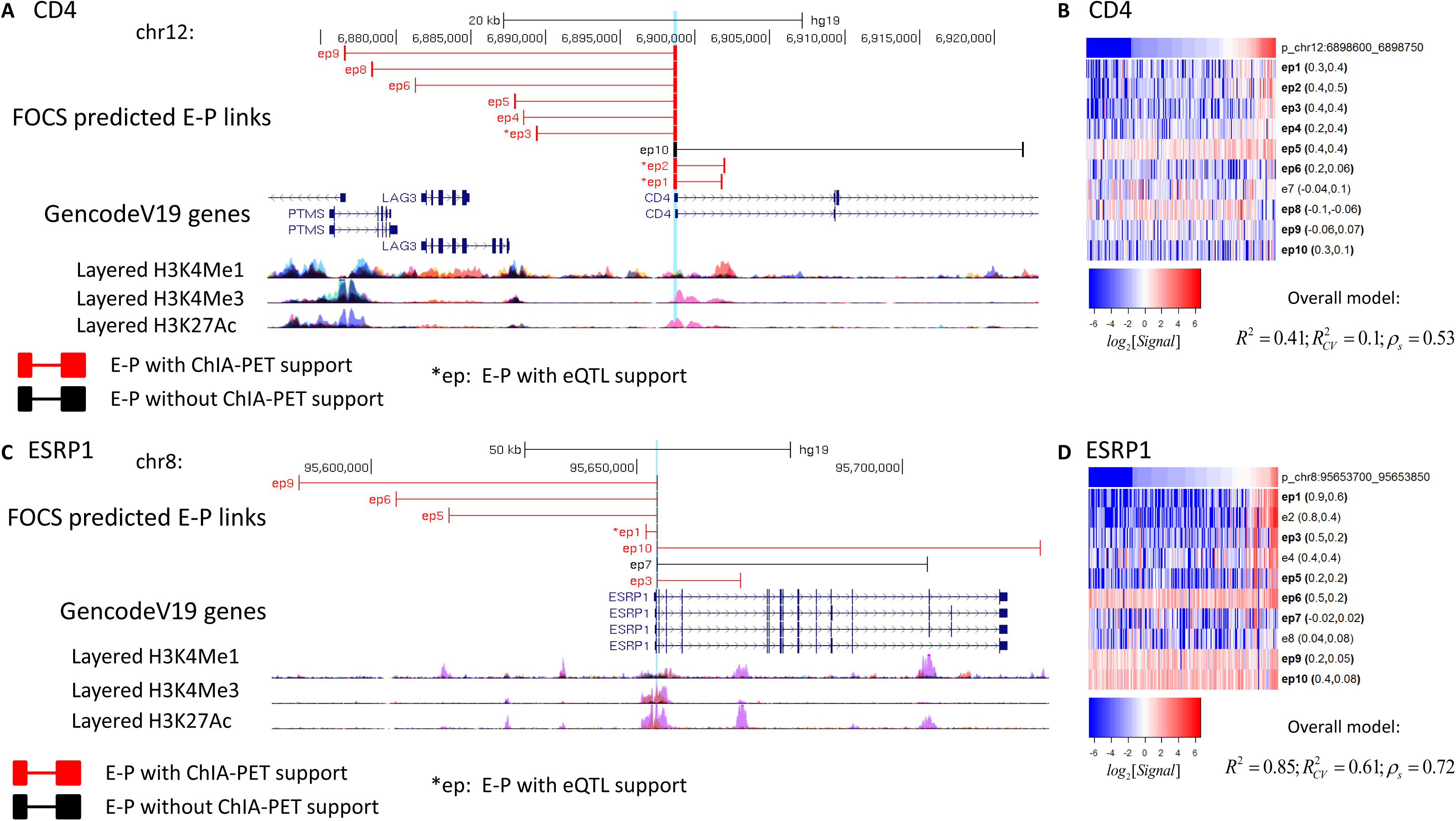
Examples of FOCS predicted E-P links supported by ChIA-PET/eQTL data. (A-B) CD4. (C-D) ESRP1. TSS location is highlighted in light blue. (B,D) Heatmaps (log_2_[RPKM Signal]) for the activity patterns of CD4/ESRP1 promoters and their 10 nearest enhancers. Enhancers included in the shrunken model are denoted by **‘ep’** and those that are not are denoted by 'e'. For each enhancer, its Pearson and Spearman correlations with the promoter are reported (left and right values in the parentheses). For each model, the *R*^2^, 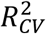, and the Spearman correlation after CV (*ρ*_*s*_) are listed.

### FOCS performance on additional large-scale datasets

Having demonstrated FOCS proficiency in predicting E-P links on the ENCODE DHS data, we next wished to expand the scope of our E-P mapping. We therefore applied FOCS to three additional large-scale genomic datasets: (1) DHS profiles measured by the *Roadmap Epigenomics* project, consisting of 350 samples from 73 different cell types and tissues; and (2) FANTOM5 CAGE data that measured expression profiles in more than 600 human cell lines and primary cells. The analysis of FANTOM5 data uses eRNA and TSS expression levels for estimating the activity of enhancers and promoters, respectively (**Online Methods**). (3) a GRO-seq compendium that we compiled. Building on eRNAs as quantitative markers of enhancer activity and the effectiveness of the GRO-seq technique in detecting eRNA expression [23], we compiled a large compendium of eRNA and gene expression profiles from publicly available GRO-seq datasets, spanning a total of 245 samples measured on 23 different human cell lines (**Online Methods**).

We applied to these datasets the same procedure that we applied above to the ENCODE data. In the analysis of these datasets, OLS yielded more validated models than the other regression methods on the Roadmap Epigenomics and GRO-seq datasets (as was the case in the ENCODE DHS data (**Fig. 1A-B**)), while GLM.NB and ZINB produced more models on FANTOM5 (**Supplementary Fig. 2A-C; Supplementary Table 1**). The performance of GLM.NB and ZINB on the FANTOM5 dataset is probably due to the high fraction of zeros entries in the count matrix of this dataset (^~^54%) compared to ENCODE, Roadmap, and GRO-seq data matrices (8%, 4%, and 19%, respectively). As OLS performed better on most datasets, all the results reported below are based on OLS. The number of promoter models that passed each validation test in each dataset is provided in **Supplementary Fig. 3A-C**. The effect of CV is presented in **Supplementary Fig. 4A-C**. In these datasets too, many of the models with high coefficient of determination (*R*^2^ ≥ 0.5) when trained on all samples, had low predictive power on novel samples 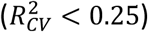 (Empirical FDR 16%, 20%, and 22% in Roadmap, FANTOM5, and GRO-seq, respectively; **Supplementary Fig. 4**), demonstrating the utility of CV in alleviating overfitting and thus reducing false positive models.

We next examined the relative contribution of each of the 10 participating enhancers to the validated models, and in these datasets too, the most proximal enhancers had the highest role, but more distal ones had very similar contribution (**Supplementary Fig. 5A**). In terms of explained fraction of the observed variability in promoter activity, 41% and 84% of the models that passed both tests in the Roadmap Epigenomics and GRO-seq datasets, respectively, had *R*^2^ ≥ 0.5, but only 11% of the validated models reached this performance in the FANTOM5 dataset (**Supplementary Fig. 5B**), probably due to its exceptionally sparse data matrix. Last, FOCS applied enet model shrinkage to the models that passed the validation tests. (The number of validated models and E-P links derived by FOCS on each dataset is summarized in **Supplementary Table 2**). In the optimally-reduced models, each promoter was linked, on average, to 3.2, 2.8 and 3.6 enhancers, in the Roadmap, FANTOM5 and GRO-seq datasets, respectively (**Supplementary Fig. 6A**), and inclusion rate decreased with E-P distance (**Supplementary Fig. 6B**). Finally, benchmarking against ChIA-PET and eQTL data, in these datasets too, FOCS outperformed the alternative methods for E-P mapping, by yielding many more E-P predictions at similar external validation rates (**Supplementary Fig. 7; Supplementary Table 3**). Collectively, we provide a rich resource of predicted E-P mapping that covers 16,349 known genes, 113,653 promoters, 181,236 enhancers, and 302,050 cross-validated E-P links.

## Discussion

In this study we present *FOCS* - a novel statistical framework for predicting E-P interactions based on activity patterns derived from large-scale omic datasets. Applying FOCS to four different genomic data sources, we derived an extensive resource of statistically cross-validated E-P links. Our E-P mapping resource further illuminates different facets of transcriptional regulation. First, a common naïve practice is to map enhancers to their nearest promoters. In FOCS predicted E-P links, ^~^26% of the enhancers are mapped to a promoter that is not the closest one (**Supplementary Fig. 8**). Second, intronic enhancers are very common - 70% of the predicted E-P links involve an intronic enhancer (**Supplementary Table 2**). Third, while on average, in the shrunken models, each promoter was linked to ^~^3 enhancers, many promoters were linked to a single dominant enhancer and some were linked to a very high number of enhancers (8-10).

As an initial step in exploring relationships between the architecture of E-P interactions and gene function, we examined the set of housekeeping genes taken from [24]. These genes are ubiquitously expressed across different cell types, suggesting that they are likely to have a simple regulation logic. Indeed, the promoters of these genes were involved in significantly lower number of E-P links compared to all other genes (p-value<0.001 in all data types; **Supplementary Fig. 9**).

We also observed that while the vast majority (^~^90%) of enhancers in FOCS-derived models had positive Pearson and Spearman correlation with the activity pattern of their target promoters, the models also included cases of negative correlation, suggesting that the regulatory element functions as a repressor (**Supplementary Fig. 10**). Finally, the activity level test in FOCS, computed using the Spearman correlation, can also account for promoter models where the relationship between the enhancer and promoter activity patterns is not linear, perhaps explaining the *R*^2^ < 0.5 values observed in the majority of FANTOM5 and Roadmap models (Fig. S5B).

One aspect that we did not consider in our analysis is the constraints imposed on transcriptional regulation by the 3D organization of the genome. Recent findings indicate that most E-P interactions are limited by chromosomal territories called topologically associated domains [25,26]. Further research is needed to better elucidate this connection between 3D organization and E-P links and to better understand to what extent such constraints are universally or differentially imposed in different cell types.

Our broad compendium of E-P interactions can greatly assist the functional interpretation of genetic variants that are associated with disease susceptibility, as the majority of such variants (^~^90%), as detected by GWAS studies, are located in noncoding sequences [27]. Similarly, it can help the interpretation of recurrent noncoding somatic mutations (SM) in cancer genomes. SM hot-spots in regulatory regions are detected at an accelerated pace with the rapid accumulation of whole-genome sequencing (WGS) of tumor samples [28,29]. Additionally, the predicted E-P links can be integrated into and boost bioinformatics pipelines that seek DNA motifs in regulatory elements that putatively regulate sets of co-expressed genes. Overall, the FOCS method that we developed and the compendium we provide hold promise for advancing our understanding of the noncoding regulatory genome.

## Methods

Methods and any associated references are available in the online version of the paper.

## Acknowledgments

T.A.H. and D.A. were supported in part by fellowships from the Edmond J. Safra Center for Bioinformatics at Tel-Aviv University. R.S. is supported by the Israeli Science Foundation (Grant 317/13) and the Dotan Hemato-Oncology Research Center at Tel Aviv University. R.E. is supported by the Israeli Cancer Association, with the generous assistance of the ICA Netherlands friends. R. E. is a Faculty Fellow of the Edmond J. Safra Center for Bioinformatics at Tel Aviv University.

## Contributions

T.A.H., R.E., and R.S. designed the research. T.A.H. and D.A. developed the computational method. T.A.H. performed the analyses, created the visualizations, parsed the ENCODE, Roadmap, and FANTOM5 data, and assembled the GRO-seq compendium. All authors analyzed the data and wrote the manuscript.

## Materials & Correspondence

Materials (code and data) are available at http://acgt.cs.tau.ac.il/focs. Correspondence should be addressed to R.E. or R.S.

## References

1. Consortium EP, others. An integrated encyclopedia of DNA elements in the human genome. Nature. Nature Publishing Group; 2012;489:57–74.

2. Thurman RE, Rynes E, Humbert R, Vierstra J, Maurano MT, Haugen E, et al. The accessible chromatin landscape of the human genome. Nature. 2012;489:75–82.

3. Consortium RE, Kundaje A, Meuleman W, Ernst J, Bilenky M, Yen A, et al. Integrative analysis of 111 reference human epigenomes. Nature [Internet]. 2015;518:317–30. Available from: http://www.ncbi.nlm.nih.gov/pubmed/25693563

4. Andersson R, Gebhard C, Miguel-Escalada I, Hoof I, Bornholdt J, Boyd M, et al. An atlas of active enhancers across human cell types and tissues. Nature. 2014;507:455–61.

5. Kim T-K, Hemberg M, Gray JM, Costa AM, Bear DM, Wu J, et al. Widespread transcription at neuronal activity-regulated enhancers. Nature. 2010;465:182–7.

6. Shlyueva D, Stampfel G, Stark A. Transcriptional enhancers: from properties to genome-wide predictions. Nat. Rev. Genet. [Internet]. Nature Publishing Group; 2014 [cited 2014 Jul 9];15:272–86. Available from: http://www.ncbi.nlm.nih.gov/pubmed/24614317

7. Elkon R, Agami R. Characterization of noncoding regulatory DNA in the human genome. Nat. Biotechnol. 2017;35:732.

8. Hah N, Danko CG, Core L, Waterfall JJ, Siepel A, Lis JT. Resource A Rapid, Extensive, and Transient Transcriptional Response to Estrogen Signaling in Breast Cancer Cells. Cell [Internet]. Elsevier Inc.; 2011;145:622–34. Available from: http://dx.doi.org/10.1016/j.cell.2011.03.042

9. Hah N, Murakami S, Nagari A, Danko CG, Kraus WL. Enhancer transcripts mark active estrogen receptor binding sites. 2013;1210–23.

10. Li W, Notani D, Ma Q, Tanasa B, Nunez E, Chen AY, et al. Functional importance of eRNAs for estrogen-dependent transcriptional activation events. Nature. NIH Public Access; 2013;498:516.

11. Melo CA, Drost J, Wijchers PJ, van de Werken H, de Wit E, Vrielink JAFO, et al. eRNAs are required for p53-dependent enhancer activity and gene transcription. Mol. Cell. Elsevier; 2013;49:524–35.

12. Andersson R, Sandelin A, Danko CG. A unified architecture of transcriptional regulatory elements. Trends Genet. Elsevier; 2015;31:426–33.

13. Léveillé N, Melo CA, Rooijers K, D'\iaz-Lagares A, Melo SA, Korkmaz G, et al. Genome-wide profiling of p53-regulated enhancer RNAs uncovers a subset of enhancers controlled by a lncRNA. Nat. Commun. Nature Research; 2015;6.

14. Core LJ, Waterfall JJ, Lis JT. Nascent RNA sequencing reveals widespread pausing and divergent initiation at human promoters. Science (80-.). 2008;322:1845–8.

15. Shiraki T, Kondo S, Katayama S, Waki K, Kasukawa T, Kawaji H, et al. Cap analysis gene expression for high-throughput analysis of transcriptional starting point and identification of promoter usage. Proc. Natl. Acad. Sci. National Acad Sciences; 2003;100:15776–81.

16. Wu H, Nord AS, Akiyama JA, Shoukry M, Afzal V, Rubin EM, et al. Tissue-specific RNA expression marks distant-acting developmental enhancers. PLoS Genet. Public Library of Science; 2014;10:e1004610.

17. Lawless JF. Negative binomial and mixed Poisson regression. Can. J. Statisitcs. 1987;15:209–25.

18. Greene WH. Accounting for excess zeros and sample selection in Poisson and negative binomial regression models. 1994;

19. Benjamini Y, Yekutieli D. The control of the false discovery rate in multiple testing under dependency. Ann. Stat. JSTOR; 2001;1165–88.

20. Benjamini Y, Hochberg Y. Controlling the false discovery rate: a practical and powerful approach to multiple testing. J. R. Stat. Soc. Ser. B. JSTOR; 1995;289–300.

21. Xie X, Ma W, Songyang Z, Luo Z, Huang J, Dai Z, et al. CCSI: a database providing chromatin‐‐chromatin spatial interaction information. Database. Oxford University Press; 2016;2016:bav124.

22. Consortium Gte, others. The Genotype-Tissue Expression (GTEx) pilot analysis: Multitissue gene regulation in humans. Science (80-.). 2015;348:648–60.

23. Danko CG, Hyland SL, Core LJ, Martins AL, Waters CT, Lee HW, et al. Identification of active transcriptional regulatory elements from GRO-seq data. Nat. Methods. Nature Research; 2015;12:433–8.

24. Eisenberg E, Levanon EY. Human housekeeping genes, revisited. Trends Genet. [Internet]. Elsevier Ltd; 2013;29:569–74. Available from: http://dx.doi.org/10.1016/j.tig.2013.05.010

25. Rao SSP, Huntley MH, Durand NC, Stamenova EK, Bochkov ID, Robinson JT, et al. A 3D Map of the Human Genome at Kilobase Resolution Reveals Principles of Chromatin Looping. Cell [Internet]. Elsevier Inc.; 2014 [cited 2014 Dec 11];159:1665–80. Available from: http://linkinghub.elsevier.com/retrieve/pii/S0092867414014974

26. Tang Z, Luo OJ, Li X, Zheng M, Zhu JJ, Szalaj P, et al. CTCF-mediated human 3D genome architecture reveals chromatin topology for transcription. Cell. Elsevier; 2015;163:1611–27.

27. Maurano MT, Humbert R, Rynes E, Thurman RE, Haugen E, Wang H, et al. Systematic localization of common disease-associated variation in regulatory DNA. Science (80-.). American Association for the Advancement of Science; 2012;337:1190–5.

28. Melton C, Reuter JA, Spacek DV, Snyder M. Recurrent somatic mutations in regulatory regions of human cancer genomes. Nat. Genet. Nature Research; 2015;47:710–6.

29. Weinhold N, Jacobsen A, Schultz N, Sander C, Lee W. Genome-wide analysis of noncoding regulatory mutations in cancer. Nat. Genet. Nature Research; 2014;46:1160–5.

